# Loop extrusion-mediated plasmid DNA cleavage by the bacterial SMC Wadjet complex

**DOI:** 10.1101/2024.02.17.580791

**Authors:** Biswajit Pradhan, Amar Deep, Jessica König, Martin Dieter Baaske, Kevin D. Corbett, Eugene Kim

**Affiliations:** Max Planck Institute of Biophysics, 60438 Frankfurt am Main, Germany; Department of Cellular and Molecular Medicine, University of California San Diego, La Jolla CA, USA; Department of Molecular Biology, University of California San Diego, La Jolla CA, USA

## Abstract

Structural maintenance of chromosomes (SMC) protein complexes play pivotal roles in genome organization and maintenance across all domains of life. In prokaryotes, SMC family Wadjet complexes structurally resemble the widespread Muk-BEF genome-organizing complexes but serve a defensive role by inhibiting plasmid transformation. We previously showed that Wadjet specifically cleaves circular DNA; however, the molecular mechanism underlying DNA substrate recognition remains unclear. Here, we use *in vitro* single-molecule imaging to directly visualize DNA loop extrusion and plasmid cleavage by Wadjet. We find that Wadjet is a symmetric DNA loop extruder that simultaneously reels in DNA from both sides of a growing loop and that this activity requires a dimeric JetABC supercomplex containing two dimers of the JetC motor subunit. On surface-anchored plasmid DNAs, Wadjet extrudes the full length of a 44 kilobase pair plasmid, stalls, and then cleaves DNA. Our findings reveal the role of loop extrusion in the specific recognition and elimination of plasmids by Wadjet, and establish loop extrusion as an evolutionarily conserved mechanism among SMC complexes across kingdoms of life.

## Introduction

SMC (structural maintenance of chromosomes) complexes are an evolutionarily conserved family of motor proteins that organize and maintain genomes through ATP-powered DNA loop extrusion. In eukaryotes, three classes of SMC complexes contribute to chromosome organization in interphase cells (cohesin), chromosome compaction in mitotic cells (condensin), and diverse DNA repair/maintenance pathways (Smc5/6) (1–3). Smc5/6 also contributes to antiviral immunity (4–8), though the mechanisms of this activity are not known. For all three eukaryotic SMC complexes, DNA loop extrusion has been directly observed in vitro (9–15). In prokaryotes, widespread SMC-ScpAB and MukBEF complexes contribute to genome organization and chromosome segregation during cell division (16–19). Some bacteria and archaea encode an SMC complex called Wadjet (also referred to as MksBEFG or EptABCD) that suppresses plasmid DNA transformation by specifically recognizing and cleaving circular DNAs less than ∼50-80 kilobase pairs (kbp) in length (20–25). While Wadjet and other prokaryotic SMCs are assumed to function by DNA loop extrusion, this activity has not been directly demonstrated.

The four-subunit Wadjet complex comprises a MukBEF-like subcomplex termed JetABC, and a fourth nuclease subunit (JetD) that forms a homodimer and is related to toprim (topoisomerase-primase) domain containing nucleases (Figure 1A). JetABC forms a pentameric complex (JetA_1_B_2_C_2_) with two copies of the SMC motor subunit JetC, two copies of the KITE (kleisin interacting tandem winged-helix elements of SMC complexes) protein JetB, and one copy of the kleisin subunit JetA. Like the related MukBEF complex, JetA_1_B_2_C_2_ complexes dimerize through their kleisin subunit JetA to yield a full complex (JetA_2_B_4_C_4_) termed a JetABC dimer. Recent studies (21, 23, 26) showed that Wadjet can restrict circular DNA through cleavage in a sequence-independent manner, sensing the topology of its bound substrate DNA through an active ATP-dependent mechanism possibly involving loop extrusion and/or translocation (27). The structural similarity of JetABC to Muk-BEF and eukaryotic SMC complexes further support the possible relation between DNA loop extrusion and plasmid restriction. However, it remains unclear whether Wadjet can extrude DNA loops or by which mechanism such motor activity would enable selective recognition and restriction of plasmid DNA.

**Figure 1.**
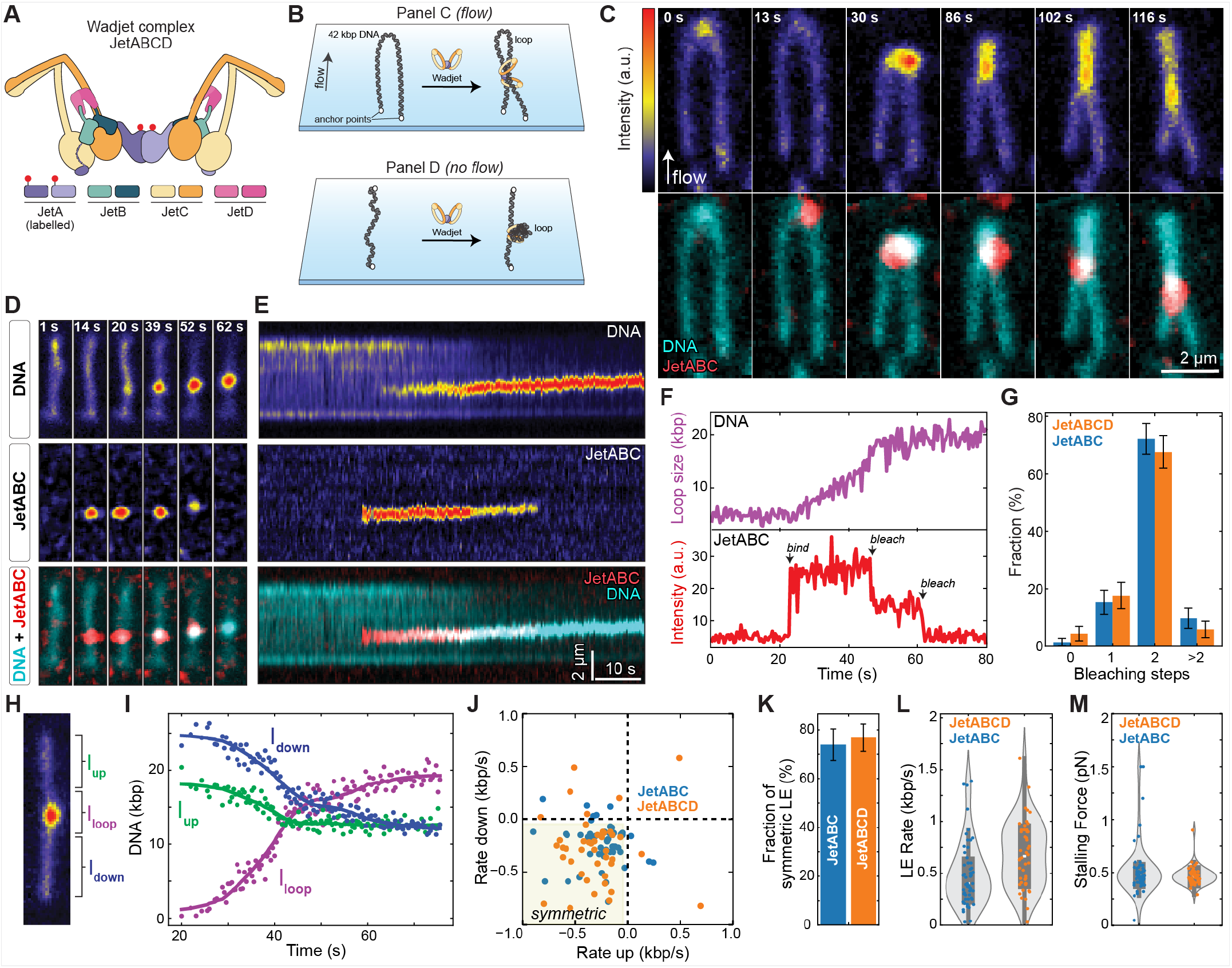
Dimers of Wadjet SMC complexes extrude DNA loops symmetrically. **(A)** Cartoon of the *P. aeruginosa* PA14 Wadjet (JetABCD) complex. **(B)** Schematics of DNA loop extrusion assays. **(C)** Series of SxO-stained DNA snapshots (top) and overlays (bottom) of DNA (cyan) and JF646-labeled Wadjet (red) showing DNA loop extrusion intermediates under constant buffer flow. **(D)** Snapshots of DNA, JetABC, their overlay (top, middle, bottom) and **(E)** the corresponding fluorescence intensity kymographs showing DNA loop extrusion in the absence of buffer flow. **(F)** Time traces of DNA loop length (top) and JetABC fluorescence intensity (bottom) determined from the kymographs in (E). **(G)** Fraction of loop-extruding Wadjet complexes that displayed either no, one, two or more bleaching steps. N=72 for JetABC and N=68 for JetABCD. **(H)** A snapshot of the looped DNA with three different regions (*I*_*loop*_, *I*_*up*_, *and I*_*down*_) indicated. **(I)** DNA lengths calculated from the kymograph in (E) for regions outside the loop (*I*_*up*_ and *I*_*down*_) and the loop region itself (*I*_*loop*_). **(J)** Rate of DNA length change for regions outside the loop determined from the linear fit of *I*_*up*_ and *I*_*down*_ at initial loop growth phase. The yellow shaded area is classified as symmetric loop extrusion. **(K)** Fraction of loop extrusion events exhibiting symmetric DNA-reeling as determined in (J). N=57 for JetABC, N=48 for JetABCD. Violin plots of Wadjet-dependent loop extrusion showing **(L)** loop growth rates and **(M)** stalling forces. N=56 for JetABC, N=49 for JetABCD.

## Results

### The bacterial SMC-family Wadjet complex extrudes DNA loops

Here, we examine DNA loop extrusion and plasmid cleavage activity of Wadjet complexes using in vitro single-molecule experiments. To this end, we separately reconstituted wild-type (WT) JetA_2_B_4_C_4_ and JetD_2_ subcomplexes from *P. aeruginosa* PA14 (21), which when mixed gave a full dimeric JetABC complex bound to two JetD dimers (JetA_2_B_4_C_4_D_4_; Figure 1A and Supplementary Figure 1). We engineered JetA to remove two native cysteine residues, and insert a cysteine residue into a disordered loop for single-site labeling with maleimide-linked JaneliaFlour 646 (JF646; Methods, Supplementary Figure 1A). The reconstituted complexes (both in the presence and absence of JetD) showed ATP hydrolysis (∼35 ATP/min/complex prior to JF646 labeling and ∼25 ATP/min/complex after labeling; Supplementary Figure 1C) that is 2 to 4-fold lower than other eukaryotic loop-extruding SMC complexes (9, 10, 15).

We tested the activity of Wadjet using a single molecule assay that allows for direct visualization of DNA loop extrusion (15). Linear biotinylated 42 kbp DNA molecules were tethered at both ends onto a passivated glass surface (Figure 1B). DNAs were stained with Sytox Orange (SxO) intercalating dye and were imaged using highly inclined and laminated optical sheet microscopy. Upon addition of Wadjet complexes with single fluorophore-labeled JetA subunits in the presence of ATP and under constant buffer flow, we observed that Wadjet binding near the midpoint of a double-tethered DNA initially resulted in the concentration of DNA into one spot at the bound Wadjet complex. Over time, the bound Wadjet moved toward the two tethered DNA ends, while the DNA elongated into a growing loop (Figure 1C and Supplementary Movie 1). No looping events by Wadjet were observed in the absence of ATP or in the presence of a non-hydrolysable ATP analogue (AMP-PNP), thereby confirming that the observed activity represents an active extrusion process (Supplementary Figure 2A).

The Wadjet-related MukBEF complex has been proposed to extrude DNA loops only when it is assembled into a dimeric complex (MukB_4_E_4_F_2_) (19, 29). To determine the stoichiometry of DNA loop-extruding Wadjet complexes, we monitored the fluorescence intensity of labeled Wadjet complexes (labeling efficiency of 70 ± 10%, Methods) and DNA during loop extrusion (Figure 1D-E, Supplementary Figure 2B, and Supplementary Movie 2-3). This analysis revealed that the Wadjet signal on DNA first increased in a single step, followed by DNA loop growth, and finally decreased in two consecutive bleaching steps (Figure 1F). Given that each JetA subunit was singly labeled, these data indicate that Wadjet dimers containing two JetA subunits are the active loop-extruding form of the enzyme. Furthermore, the quantification of the observed ratio of zero (unlabelled), one and two bleaching steps during looping events (Figure 1G) showed the majority events (72% for JetABC and 67% for JetABCD) were two bleaching steps and the ratios agreed with the expected ratio of Wadjet dimers to monomers (Supplementary Figure 2C), indicating that the majority of loop-extrusion events are performed by Wadjet dimers. To test whether these dimers are comprised of two motors simultaneously reeling in DNA from both sides, i.e. “symmetric” loop extrusion, using the terminology established by Barth et. al. (30), we estimated the length of DNA within the loop and on each side of the loop during the initial phase of loop growth (30 s, Figure 1H-I, Supplementary Figure 2D-E). We observed that DNA from both sides of the loop are simultaneously reeled in during loop growth (Figure 1I), and that the reeling rates from each side are often different (Figure 1J), suggesting the action of two independent, but physically linked, motors. The majority of the observed looping events occurred in a symmetric fashion (78 ± 6% for JetABC and 74 ± 6% for JetABCD, Figure 1K), with the average growth rates of 0.5 ± 0.3 kbp/s (for JetABC) and 0.7 ± 0.3 kbp/s (for JetABCD, Figure 1L), a value 2 to 3-fold lower than those reported for eukaryotic SMC complexes (1-1.5 kbp/s) (9, 10, 15). The average stalling forces, estimated from the value of relative DNA extension at which loop extrusion was halted and converted to the known force–extension relation (Supplementary Figure 2F-G) (10), were 0.5 ± 0.2 pN (for JetABC) and 0.5 ± 0.1 pN (for JetABCD, Figure 1L), values similar to the stalling forces measured for eukaryotic SMC complexes (0.2-0.5 pN). However, the comparably slower extrusion process mediated by Wadjet dimer led to stable loops with little or no loop diffusion or slippage (Supplementary Figure 2H). Notably, we found that the presence of JetD did not influence the general characteristics of loop extrusion mediated by JetABC (Figure 1G and 1J-M).

### Wadjet dimerization is essential for DNA loop extrusion

To directly test whether Wadjet dimerization is necessary for DNA loop extrusion, we designed two artificial monomeric constructs, M1 and M2, by truncating the kleisin subunit JetA. We used a differential tagging strategy to purify complexes with one full-length JetA subunit and one JetA subunit lacking its C-terminal JetC-binding winged-helix domain (M2; yielding JetA_2_B_4_C_2_) or missing both the winged-helix domain and JetB-binding disordered region (M1; yielding JetA_2_B_2_C_2_) (Materials and Methods, Figure 2A and Supplementary Figure). In both M1 and M2, only the single full-length JetA subunit is fluorophore-labeled. Mass photometry (Figure 2B) confirmed that both M1 and M2 complexes are primarily monomeric, while a minor population (3% for M1 and 5% for M2) are dimeric and presumably contain two full-length JetA subunits. Single-molecule DNA loop extrusion assays with these complexes revealed a ∼15-fold reduction in looping probability compared to WT (75% for WT, 4% for M1 5% for M2; Figure 2C) and thus in agreement with the model that dimerization is required for loop extrusion mediated by Wadjet. The overall DNA binding probabilities of these mutants were also significantly lower than for WT (Figure 2D). The rare loop extrusion events that were observed with M1 and M2 proceeded symmetrically (Figure 2F, 2H, 2J-K and Supplementary Figure 4A-B), suggesting that loop extrusion is mediated by the dimeric subpopulation in each construct (Figure 2B). This hypothesis is further supported by our finding that loop-extruding Wadjet complexes for both M1 and M2 predominantly exhibit two-step photobleaching (Figure 2E, 2G and 2I) with statistics matching a dimer model closer than a monomer model (Supplementary Figure 4C), indicating that these complexes contain two full-length JetA subunits and are therefore dimeric. The average extrusion rate (0.7 ± 0.4 kbp/s for M1, 0.7 ± 0.3 kbp/s for M2; Supplementary Figure 4D) and stalling force (0.5 ± 0.2 pN for M1, 0.5 ± 0.2 pN for M2; Supplementary Figure 4E) were also similar to the values obtained for WT (Figure 2L-M). The comparative analysis of the loop formation probability of WT, M1 and M2 per DNA loading (Figure 2L) showed that for WT, most DNA binding events led to loop extrusion (82%), while more non-looping events, either DNA binding followed by dissociation (Figure 2M) or translocation (Figure 2N), were observed for M1 and M2 (79% and 64% of non-looping events for M1, M2; Figure 2L). Altogether, these data strongly support a model in which the DNA loop-extruding form of Wadjet is a dimer.

**Figure 2.**
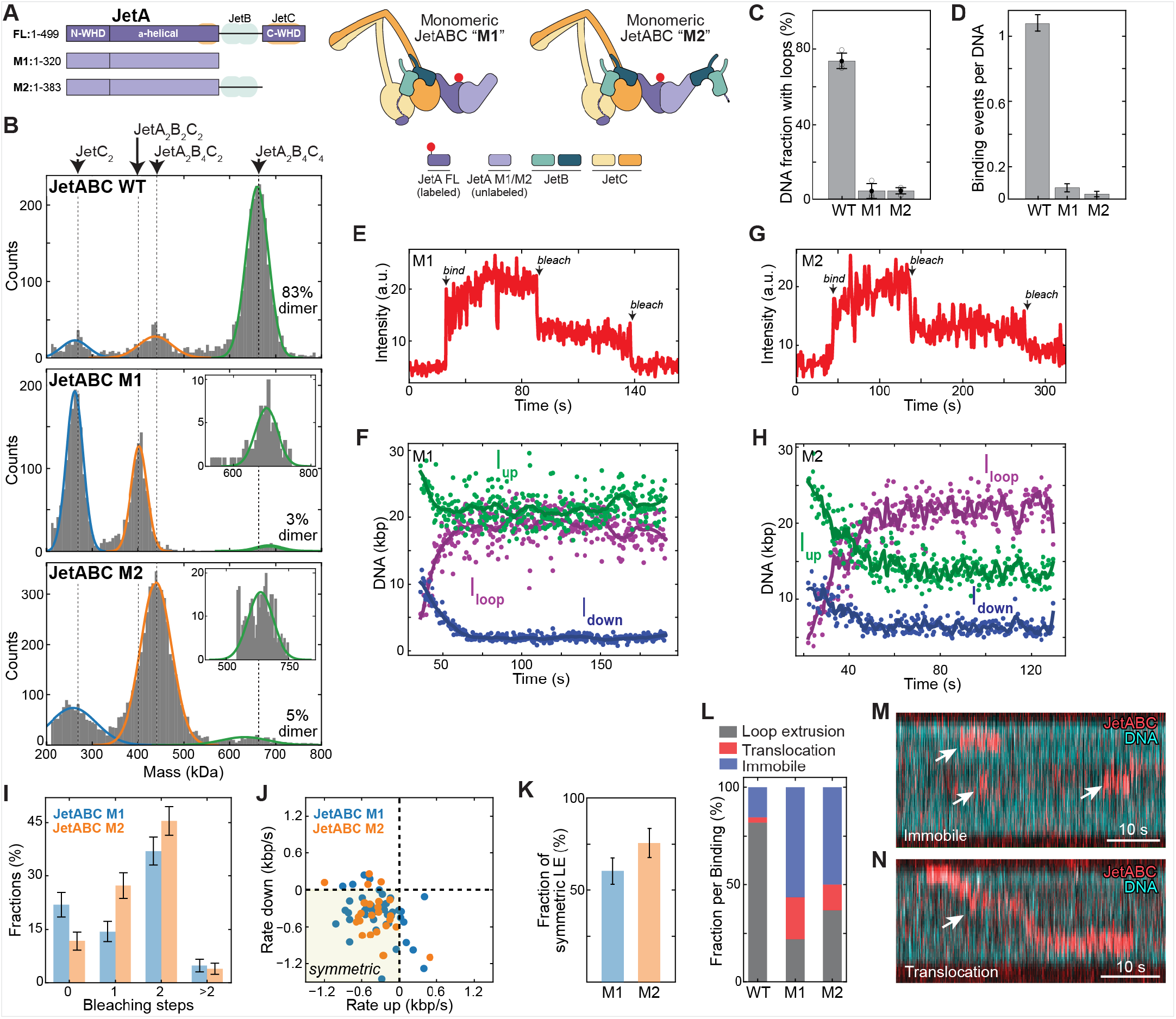
Dimerization is essential for Wadjet-mediated loop extrusion. **(A)** Cartoon of the designed two truncated monomeric constructs, named as “M1” (JetA_2_B_2_C_2_) and “M2” (JetA_2_B_4_C_2_). **(B)** Histograms of mass distribution for wild-type (WT; top), M1 (middle) and M2 (bottom). Inset: zoomed-in peak centred around 670 kDa, indicating the respective dimer fractions. For fitting the dimer peak of M2, the mass values lower than 540 kDa were excluded. **(C)** Fraction of DNA molecules that formed loops following the addition of WT, M1 and M2 Wadjet complexes. N=318, 467, 459 for WT, M1, M2, respectively. **(D)** Probabilities of Wadjet/DNA binding per DNA molecule for WT, M1, M2. N=101 for each WT, M1, M2. **(E, G)** Time traces of Wadjet fluorescence intensity and **(F, H)** the corresponding time traces of DNA lengths during DNA looping for M1 (E, F) and M2 (G, H). (l)Fraction of loop-extruding M1 and M2 complexes that displayed either no, one, two or more bleaching steps. N=146 for M1, 154 for M2. **(J)** Scatter plot showing the rates of DNA length change for regions outside the loop. The yellow shaded area is classified as symmetric loop extrusion. **(K)** Fraction of symmetric loop extrusion events for M1 (N=44) and M2 (N=26) **(L)** Fraction of looping, immobile and translocating events upon DNA binding for WT, M1 and M2; N=101 each for WT, M1, M2. **(M)** Example kymograph of immobile Wadjet complexes bound on DNA. **(N)** Example kymograph of a translocating Wadjet complex on DNA.

### Wadjet stalls at the plasmid anchor point after extrusion of a full-length DNA circle

Next, we investigated the relation of loop extrusion and plasmid cleavage activity of Wadjet. Since Wadjet specifically cleaves circular DNAs up to 50-80 kbp in length (23), we tethered circular 44-kbp DNAs to a glass surface at a single point and stained with Sytox Orange (Materials and Methods, Figure 3A). Upon application of buffer flow, the surface-anchored DNAs are stretched to extents slightly less than half their length, and exhibited slightly more than twice the fluorescence intensity of an equivalent-length linear DNA (Figure 3B). Using JetABC in the absence of JetD, we first monitored loop extrusion on these flow-stretched plasmid substrates (Figure 3C-3F). Upon binding, Wadjet complexes quickly approached the anchor point of the surface-tethered plasmid (Figure 3C-3F). When Wadjet initially binds in the middle of a stretched plasmid, the movement of the complex is accompanied by folding/compaction of the plasmid into two segments that correspond to extruded and non-extruded segments of the plasmid, until Wadjet reaches the tether point. Continued reeling-in of the unextruded segment after encountering the tether then leads to progressive unfolding/decompaction as Wadjet eventually extrudes the full length of the plasmid (Figure 3C-D). On the other hand, when Wadjet initially binds near the tip of the plasmid DNA, its DNA reeling-in action barely changes the overall extent of the stretched plasmid and the labeled Wadjet complex appears to simply translocate along the length of the stretched DNA (Figure 3E-F); this behavior would be expected from a symmetric loop-extruding motor. Quantification of the photobleaching statistics again confirmed the dimeric nature of Wadjet complexes bound to plasmid DNAs (Figure 3G). Notably, the majority of loop-extruding Wadjet dimers initially bound in the middle or near the tip of the plasmid, extruded a loop, then stalled after reaching the anchor position (Figure 3H). This stalling of Wadjet complexes led to accumulation of multiple complexes at anchor points over time (Figure 3I, Supplementary Movie 4). In contrast, when Wadjet dimers extrude loops on single-tethered linear DNAs with one free end, they either dissociate from the DNA upon encountering the free end (Figure 3J) or remain at the anchor (Supplementary Figure 5), thus leading to a relatively low number of DNA-bound complexes as compared to plasmid (Figure 3K).

**Figure 3.**
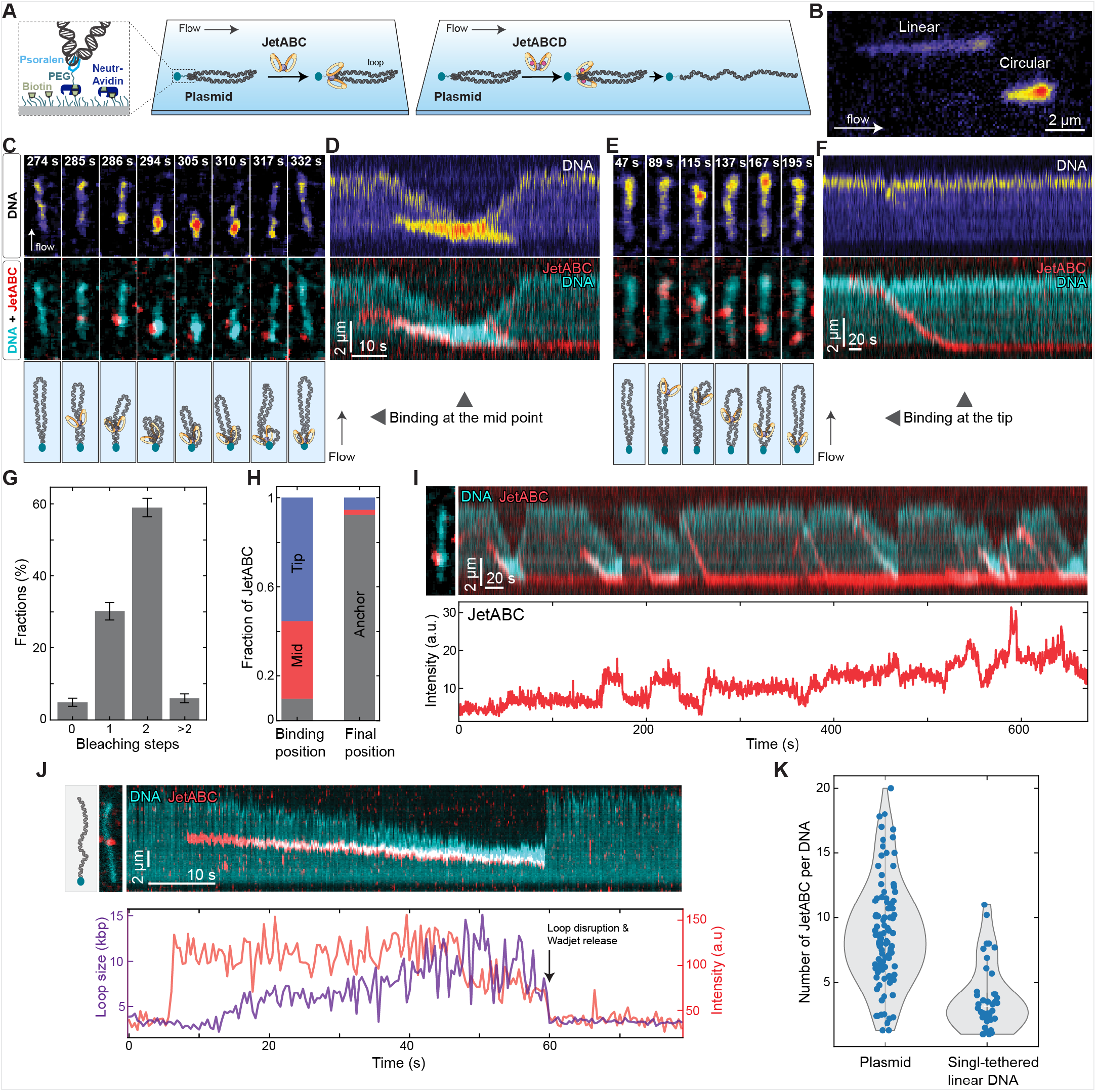
Loop-extruding Wadjet complexes stall at the anchor point of the surface-tethered plasmid. **(A)** Schematic of loop extrusion assay for surface-tethered plasmid DNA. **(B)** Snapshot showing intensity and length difference in a 44 kbp plasmid DNA tethered at one point and a 44 kbp linear DNA tethered at one end of the DNA. **(C, E)** Snapshots of DNA (top), its overlay with JetABC (middle), schematics (bottom) and **(D, F)** the corresponding fluorescence intensity kymographs showing two representative examples of loop extrusion events on plasmid DNA. **(G)** Fraction of loop-extruding JetABC complexes that displayed either no, one, two or more bleaching steps on plasmid DNA. N=368 events. **(H)** Estimation of initial and final position of loop-extruding JetABC complexes. N=112 events. **(I)** Kymograph of JetABC and DNA (top) and the corresponding time traces of JetABC fluorescence signal (bottom) showing multiple loop extrusion events leading to accumulation of Wadjet molecules at the surface anchor of plasmid. (J) Overlaid kymograph (top) and the corresponding fluorescence time traces (bottom) of JetABC (red) and DNA (cyan, purple) during loop extrusion on single-tethered linear DNA. (K) Comparison of number of JetABC complexes remained on plasmid or single-tethered linear DNA after 20 minutes of incubation, N=101 for plasmid, N=40 for single-tethered linear DNA.

### Wadjet-mediated DNA cleavage relies on a full-length extrusion of plasmid DNA

Our observation that loop-extruding JetABC complexes stall near the surface anchor point of the plasmid DNA raised the possibility that the remaining short unextruded part of the plasmid might be a favorable substrate for JetD-mediated DNA cleavage in the full complex (26). We therefore tested the activity of fully-reconstituted JetABCD complexes (see Materials and Methods) on plasmid DNA. We observed that in the presence of JetD, DNA loop extrusion was occasionally followed by DNA cleavage, as observed by an abrupt increase in the extent of stretched DNA and a corresponding reduction of DNA fluorescence intensity (Figure 4A, Supplementary Figure 6-7 and Supplementary Movie 5-7). The estimation of the end-to-end length distributions of the cleaved product (Figure 4B) indicate that JetD-mediated cleavage occurs near the anchor point (Figure 4C), where loop-extruding Wadjet complexes remain stalled (Figure 3H). Importantly, we observed that a significant fraction of plasmids (∼14%) were linearized, only in the presence of JetD and ATP, while almost no cleavage occurred (<1%) in the absence of ATP, in the absence of JetD or JetABCD, or when JetD was mutated to eliminate its nuclease activity (E365A; Figure 4D). Moreover, incubation of JetABCD complexes with linear DNA (single- or double-tethered) did not lead to DNA cleavage (Figure 4D). The small fraction of linearized DNA in all cases presumably arises from light-induced non-specific cleavage activity, as evidenced by the fact that we observe a similar fraction of cleavage without any proteins (Figure 4D). In further support of the loop-extruding Wadjet-mediated DNA cleavage, we observed that nearly all DNA cleavage events (98%) occurred after full-length extrusion of the plasmid (Figure 4E) Furthermore, ∼9% of the cleavage events occurred after only one loop extrusion event by Wadjet complex (Figure 4F-G, Supplementary Figure 7C and Supplementary Movie 7), suggesting that a single loop-extruding Wadjet dimer complex is sufficient for plasmid cleavage. Collectively, our data strongly suggest that the plasmid cleavage activity of Wadjet complex is linked to DNA loop extrusion.

**Figure 4.**
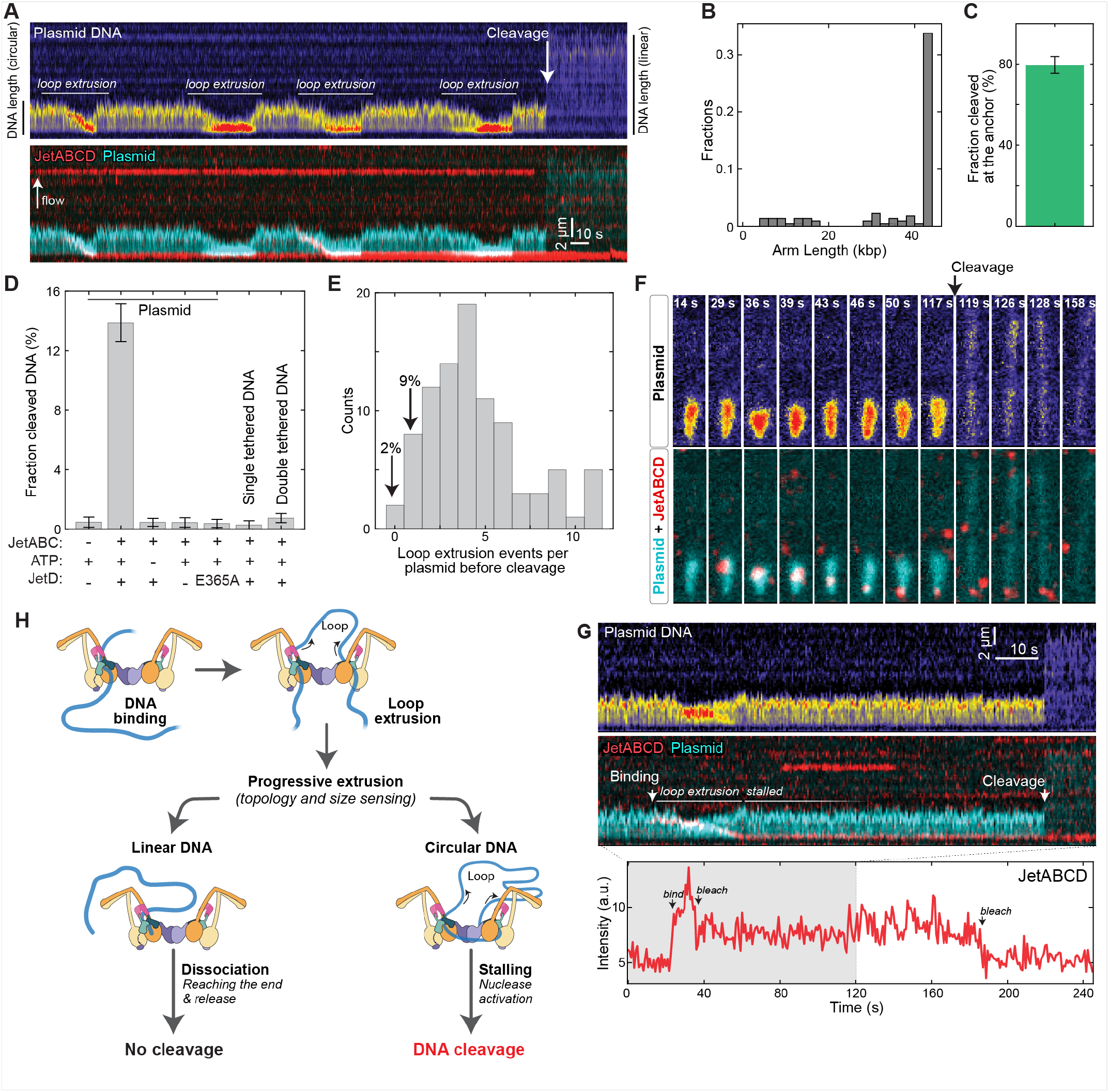
JetABCD cleaves plasmid through loop extrusion. **(A)** Example kymograph of DNA (top) and its overlay with JetABCD signal (bottom) showing multiple loop extrusion events followed by DNA cleavage. **(B)** Distributions of the end-to-end length of the flow stretched DNA after cleavage. N=101 events. **(C)** Fraction of cleavage events that occurred at the plasmid’s surface anchor point. **(D)** Fractions of plasmid cleavage events without JetABCD, in the presence of ATP and JetABCD, without ATP, without JetD, with JetABCD containing nuclease activity-deficient JetD mutation (E365A), or when using single-/double-tethered linear DNA. N_tot_ = 416, 743, 641, 437, 509, 348, and 796 molecules, respectively. **(E)** Distribution of number of looping events occurred before DNA cleavage. **(F)** Snapshots of Plasmid DNA (top) and its overlay with JetABCD (bottom), **(G)** the corresponding kymographs of DNA (top), its overlay with JetABCD (middle) and the corresponding fluorescence signal of JetABCD (bottom) showing a single JetABCD dimer-mediated loop extrusion followed by DNA cleavage. **(H)** Model showing loop extrusion-mediated DNA topology sensing and cleavage by the bacterial Wadjet complex.

## Discussion

The prokaryotic Wadjet defense system exhibits the unique ability to recognize and specifically cleave circular DNA molecules in a sequence-independent manner, thereby protecting its host from transformation by plasmids (21, 23). Prior biochemical and structural work on this complex from two different bacteria revealed structural similarity to SMC complexes, and suggested that Wadjet might couple DNA loop extrusion by the JetABC subcomplex to DNA cleavage by the JetD subunit to accomplish its defense role. Here, we show that JetABC extrudes DNA loops symmetrically, and that JetABCD can cleave circular DNAs after extruding the full length of a circular DNA substrate and stalling. Wadjet complexes also stall on double-tethered linear DNAs but do not cleave these DNAs, suggesting that stalling upon full-length extrusion of a circular DNA induces a particular conformation necessary for DNA cleavage. When combined with a recent cryo-electron microscopy structure revealing JetD poised for cleavage of a (presumably extruded) DNA substrate (26), our data strongly support a model in which (1) Dimer of JetABCD loads onto DNA and processively and symmetrically extrudes a DNA loop; (2) stalls upon extrusion of the entire length of a circular DNA and enters a distinct “stuck” conformation that (3) activates the JetD subunit to cleave DNA and allow Wadjet to dissociate (Figure 4H).

Eukaryotic loop-extruding cohesin and condensin complexes possess a single DNA-binding SMC dimer and are therefore termed “monomeric” complexes. A key element of these complexes’ loop extrusion activity comes from a HEAT-repeat containing subunit (termed HAWK) that provides an additional DNA binding site in the complex (1–3). In contrast, prokaryotic SMC complexes like MukBEF and JetABC do not possess HAWK subunits; rather, they possess KITE subunits (MukE and JetB, respectively) that are thought to help clamp DNA on their respective SMC subunits (MukB and JetC) but do not constitute an independent DNA binding site (21, 31). This structural feature of prokaryotic complexes, plus the ability of their atypical kleisin subunits (MukF and JetA) to form homodimers, led to the idea that these complexes might require two SMC dimers for DNA loop extrusion. This model is supported by data showing that disrupting dimerization of the MukF kleisin leads to phenotypes resembling MukBEF deletion mutants, including chromosome organization defects and the production of anucleate cells (19). Our data showing that Wadjet, which structurally resembles MukBEF, loads onto and extrudes DNA loops as a dimeric complex demonstrates that this model is correct. We find that the rates at which DNA is reeled in from each side by Wadjet can vary, suggesting that DNA loop extrusion is accomplished by two independently acting, but physically tethered, motors. Similarly, another KITE-based system, Smc5/6 was recently shown to be a symmetric motor that extrudes a DNA loop upon dimerization (15) indicating dimer-mediated loop extrusion might be a common feature among KITE-based SMC complexes.

In conclusion, our study unambiguously demonstrates that the bacterial Wadjet complex discerns the circular topology of plasmid DNA through its loop extrusion action. A dimeric Wadjet complex, which is a minimal functional unit for loop extrusion, reels in DNA simultaneously from both sides on both circular and linear DNAs, while DNA cleavage occurs only for the plasmid DNA, specifically at the anchor point of the plasmid DNA where loop extrusion was stalled. Our findings provide direct evidence that loop extrusion promotes selective recognition and restriction of plasmids, thus revealing its role in immune system. Furthermore, this work establishes loop extrusion as a unifying principle for genome folding shared by evolutionally conserved SMC complexes.

## Acknowledgements

The authors thank Gabriele Maul for help in DNA preparation used in this study. The authors acknowledge funding from the National Institutes of Health R35 GM144121 (to K.D.C.), Max Planck Society, and European Research Council Starting Grant 101076914 (to E.K.), and Hessian Ministry of Science and Art MSCA Grant (to B.P.). This paper was typeset with the bioRxiv word template by @Chrelli: www.github.com/chrelli/bioRxiv-word-template.

## Author contributions

Conceptualization: B.P., A.D.

Methodology: B.P., A.D., M.D.B.

Investigation: B.P., A.D., J.K.

Data curation: B.P., A.D.

Visualization: A.D., B.P., K.D.C., E.K.

Funding acquisition: K.D.C., E.K.

Project administration: K.D.C., E.K.

Supervision: K.D.C., E.K.

Writing – original draft: E.K.

Writing – review & editing: K.D.C., E.K., A.D., B.P., M.D.B.

## Data and materials availability

Original imaging data and protein expression constructs are available upon request.

## Supplementary Material

Supplementary material (Supplementary Figures 1-7 and Supplementary Movies 1-7) is available online at bioRxiv.

## Materials and Methods

### Protein construct design

To design a site-specific maleimide-labelable Wadjet complex, a previously used His6-JetA + untagged JetB coexpression construct (1) was modified to remove native cysteine residues in JetA (C36A, C355A) and insert a cysteine into a disordered loop region of JetA (C66; this residue was added between JetA residues A65 and S66 (original position) of the wild type sequence). To design monomeric JetABC complexes, we cloned N-terminally StrepII-tagged JetA (C36A, C355A) and untagged JetB into UC Berkeley Macrolab vector 13S-A (Addgene # 48323). To generate truncated JetA constructs, stop codons were introduced after residue position 320 (M1) or 383 (M2). Expression vectors for untagged JetC and His6-tagged JetD (wild type or E365A mutant) were unchanged from our prior study (1).

### Protein purification and complex reconstitution

Purification of JetABCD complexes is schematized in Supplementary Figure 1A, and purification of JetABC M1 and M2 complexes is schematized in Supplementary Figure 3A. Briefly, proteins were expressed in *E. coli* Rosetta2 pLysS (EMD Millipore) by growing cells in 2XYT media at 37°C to an OD600 of 0.55-0.75, followed by induction with 0.33 mM IPTG. Cultures were incubated overnight (∼15 hours) at 20°C for protein expression. Cells were harvested by centrifugation, and pellets were resuspended in ice-cold resuspension buffer containing 50 mM Tris pH 7.5, 300 mM NaCl, 10 mM imidazole, 10% glycerol, 2 mM β-mercaptoethanol. Resuspended cells were lysed by sonication and the lysate was clarified through centrifugation. Proteins were purified using Ni^2+^ affinity chromatography (Ni-NTA Superflow, Qiagen). To purify JetAB complexes containing one full-length and one truncated JetA protomer, vectors encoding His6-JetA (full length)+JetB and StrepII-JetA (M1 or M2)+JetB were coexpressed in the same cells, then purified using successive Ni^2+^ and Strep-Tactin (Strep-Tactin XT 4Flow) affinity chromatography. For JetC purification, tagless JetC efficiently binds Ni^2+^ affinity resin; so tagless JetC was employed for purification. After Ni^2+^ and optionally Streptactin chromatography, JetAB and JetC proteins were further subjected to anion-exchange chromatography (HiTrap Q HP, Cytiva) in a buffer with 20 mM Tris pH 7.5, 2 mM β-mercaptoethanol, and 50 mM to 1 M NaCl. Eluted proteins (JetAB, JetC, JetD, or JetDE365A) were concentrated and then passed over a Superose 6 Increase 10/300 GL size exclusion column (Cytiva) in a buffer containing 20 mM HEPES pH 7.0, 150 mM KCl, 1 mM TCEP (tris(2-carboxyethyl) phosphine). Purified JetAB (or conjugated JetAB, see below) and JetC subunit proteins were mixed in a specific stoichiometric ratio to obtain the JetA2B4C4 complex, and any aggregated particles were removed through another round of size exclusion chromatography. JetD was added to this preformed JetABC complex to reconstitute full JetA2B4C4D4 complex. Purity of samples was determined using SDS-PAGE analysis and samples were flash-frozen in liquid nitrogen and stored at -80°C until use.

### JetA labeling

A maleimide-based labeling approach was employed for fluorescent tagging of the Wadjet complex using the inserted cysteine 66 on JetA. Following the first round of size exclusion chromatography in a buffer containing 20 mM HEPES pH 7.0, 150 mM KCl, 1 mM TCEP, the JetAB subcomplex was mixed with Janelia Fluor 646 (Tocris) dye at a 1:20 molar ratio for JetAB:dye. The mixture was incubated overnight at 4°C with constant gentle rotation. On the subsequent day, excess dye was removed by passing the complex over a Superose 6 Increase 10/300 GL size exclusion column (Cytiva), and the conjugation efficiency was calculated through measurements obtained using NanoDrop (Thermo Scientific). The average labeling efficiency was determined to be ∼70% for the two labeling sites in a JetA2B4 complex. The conjugated JetAB sample was then mixed with other subunits for Wadjet full complex reconstitutions (see above).

### ATPase assays

ATP hydrolysis was measured using the ADP-Glo Kinase assay kit (Promega), following the protocol provided by the manufacturer, as described previously (1). 5 µL reactions were performed in a buffer containing 50 mM HEPES pH 7.5, 20 mM KCl, 1 mM DTT, 10 mM MgCl2, plus 1 mM ultra-pure ATP and 200 nM of Wadjet complex (JetA2B4C4 or JetA2B4C4D4). Reactions were incubated for 35 minutes at 37°C, followed by the addition of ADP Glo reagent then kinase detection reagent as specified by the manufacturer (Promega). Luminescence was measured in a 384-well plate using a TECAN Infinite M1000 microplate reader (Mannedorf, Switzerland). A standard curve was calculated using a known concentration mixture of ADP and ATP. The measurements are presented as the mean of four technical replicates with standard deviation.

### Single-molecule assays

The single molecule experiments were performed as described in a previously established protocol by Pradhan et. al. (2) with some modifications. Here we describe the details in the following sections.

### Microscopy

A custom-built highly inclined optical light sheet microscope was used for the imaging of DNA and single molecules. In brief, lasers with wavelengths 561 nm (coherent) and 638 nm (cobolt) were coupled with a single-mode optical fiber. Subsequently, the light was directed into a Zeiss microscope (AxioVert200) via a mirror with which the angle of illumination to the objective lens can be manually adjusted. The total internal reflection objective (alpha-Plan-APOCHROMAT ×100/1.46 numerical aperture, oil) in combination with off-center laser beam coupling resulted in a thin light sheet propagating through the sample. This enabled selective illumination of the DNA and proteins above the surface while reducing background intensity from fluorophores located in regions outside the light sheet. The laser reflection and scattering were separated and suppressed with a dichroic filter (no. t405/488/561/640rpc2, Chroma) in combination with a multiband notch filter (no. NF03-405/488/561/635E-25, Semrock). Fluorescence images were recorded using a sCMOS camera (PCO edge 4.2) operated by custom-developed software. For simultaneously imaging of DNA and SMC complexes, an alternative excitation was used through electronic synchronization of an acoustic-optic tunable filter (MPDSnCxx-ed1-18 and AOT-FnC_MDS driver from AA-Optoelectronic). The images were stored and visualized in napari and pyqtgraph. Typically, a measurement involved recording 10000 images with an acquisition time of 100 ms, resulting in a total measurement duration of 1000 seconds.

### Slide functionalization and flow cell preparation

Glass coverslips (1 mm thickness, 25 mm x 60 mm) were drilled with lasers to make holes on both side (20 mm separation) of it such that the holes fit to 200 µL pipette tips (#catalog). Microscope slides (#catalog, 24 mm x 60 mm) and the support coverslips were thoroughly cleaned by i) sonication in acetone, water and finally acetone over periods of 10 minutes, each; ii) sonication in 1 M KOH for 30 mins; iii) treatment with acid piranha (sulfuric acid and hydrogen peroxide, 5:1) followed by rinsing in Milli-Q water. Post-cleaning the slides/coverslips were extensively washed with methanol to remove residual moisture. Afterwards, the slides and coverslips were silanized with 5 % 3-[(2aminoethyl)aminopropyl]trimethoxysilane in methanol containing 5% glacial acetic acid with 30 min incubation time. The silanized slides and coverslips were washed with methanol and subjected to drying in an oven at 60°C for 3 hours leaving amine groups accessible on the surface. Finally, the slides and coverslips were reacted with 20 mg/ml methoxy-PEG-N-hydroxysuccinimide (no. MW 3500, Laysan Bio) and 0.2 mg/ml biotin-PEG-N-hydroxysuccinimide (no. MW3400, Laysan Bio) in 50 mM borate buffer pH 8.5 for 30 mins and then rinsed with water and dried. This last step was repeated thrice, and the slides were finally dried with compressed nitrogen and stored at -20°C until further use.

For flow cell assembly, a coverslip and a functionalized slide were sandwiched using channel patterns created from double sided adhesive tape (100 µm thickness). The channels with volume ∼10 µL were made by manually cutting the tape such that it leaves an inlet and an outlet on each side of the channel. For sideflow application, three-way channels in a µ shape were constructed. Any open ends were sealed with epoxy to prevent leakage of liquid. One side of the channel was fitted with a tube linking it to a syringe pump to facilitate controlled fluid movement. The temperature within the flow cell was maintained at 30 0C, achieved by attaching a self-adhesive heating foil (Thermo TECH Polyester Heating foil self-adhesive 12V DC, 12 V AC 17 W IP rating IPX4 (L × W) 65 × 10 mm2) to the upper surface of the coverslip.

### DNA substrates for looping and cleavage assay

The non-coilable 42-kbp linear DNA construct with biotin-handles at both ends and the 44-kbp plasmid were synthesized using cosmid-I95. The cosmid-I95 plasmid was amplified in a NEB 10-beta (New England Biolabs, C3019H), and the DNA was purified using a QIAfilter Plasmid Midi Kit (Qiagen, 12243). Biotin-containing handles were made using a PCR on pBluescript SK+ (Stratagene) with GoTaq G2 DNA polymerase (Promega, M7845). The PCR was done using biotinylated primers primer CD21 (GAC-CGAGATAGGGTTGAGTG) and CD22 (CAGGGTCGGAACAGGA-GAGC), resulting in a 1,238 base pair (bp) DNA fragment that contained one biotin. This was cleaned up using a PCR cleanup kit (Promega, A9282). The biotin handle and cosmid-I95 DNA were both digested for 2 h at 37 °C with SpeI-HF (New England Biolabs, R3133L) and subsequently heat-inactivated for 20 min at 80 °C, resulting in linear ∼42-kbp DNA and ∼600-bp biotin handles. The digested products were mixed together, and we used a 20:1 molar excess of the biotin handle to linear cosmid-I95. We then added T4 DNA ligase (New England Biolabs, M0202L) in the presence of 1 mM ATP overnight at 16 °C and subsequently heat-inactivated the next morning for 10 min at 65 °C. The resulting 42-kbp DNA construct was cleaned up using ÄKTA pure, with a homemade gel filtration column containing approximately 49 ml of Sephacryl S-1000 SF gel filtration media (Cytiva), run with TE + 150 mM NaCl2 buffer. The sample was run at 0.25 ml min−1, and we collected 0.5 ml fractions. For experiments requiring tethering of only one end of the linear DNA, the 42-kbp DNA was biotinylated at one end and a dig ligand at the other end.

### Plasmid biotinylation

2mM of biotin-Peg5000-amine was incubated with 1 mM succinimidyl-[4-(psoralen-8-yloxy)]-butyrate in DMSO overnight at RT which resulted in biotin-Peg5000-psoralen. The reaction was quenched by dilution in 40 mM Tris-HCl pH 7.5 buffer and to a final concentration of 1 µM. The purified cosmid-I95 plasmid (∼44-kbp) was incubated for 30 min at RT with 1uM biotin-Peg5000-psoralen in a ratio of 1:1 in 1x TE buffer following an incubation on ice for 1 min. The tube was then exposed to UV light (365 nm) in RT for 1 h. The sample was diluted in TE buffer to a total volume of 50 µl and dialyzed 2 times for at least 5 hours using Slide-A-Lyzer™ MINI Dialysis Unit, 10 K MWCO, 0,1 ml (Thermo, YG359049) with an exchange of the TE buffer in between. This process resulted in no more than one biotin per plasmid as observed from the number of tethering points of the plasmids on streptavidin coated slides.

### Loop extrusion and cleavage assays

In the flow cell, the biotinylated surface was incubated with 1 µM streptavidin/neutravidin for 5 min using T100 buffer (40 mM Tris-HCl pH 7.5, 100 mM NaCl), followed by washing with the same buffer. Subsequently, the double-biotinylated DNA was flown through the channels with a flow rate of 2.5 to 3 µL/min using a syringe pump and then flushed with T100 buffer with 0.5 mg/mL BSA to further prevent the nonspecific binding the SMC complexes to the surface. This resulted in the formation of double-tethered DNA (dtDNA). A similar process was applied to single biotinylated linear DNA leading to the formation of single-tethered DNA (stDNA). The density of the DNA was closely monitored through real-time visualization via intercalation with Sytox Orange (SxO). The flow of DNA was stopped once the density reached 100 to 200 DNA molecules per 100 µm x 100 µm area. The channels were then equilibrated with the imaging buffer (100 mM Tris-HCl pH 7.5, 100 mM NaCl, 7.5 mM MgCl2, 0.5 mg ml–1 BSA, 0.2 mM TCEP, 2 mM ATP, 100 nM SxO, 30 mM d-glucose, 2 mM trolox, 10 nM catalase, 37.5 µM glucose oxidase), preheated to 30°C. For side flow experiments, 200 nM SxO was used instead. The loop extrusion process was then monitored, typically using Wadjet with a concentration of 100 pM in the imaging buffer.

Plasmids (100 pM) labeled with Psoralen-Biotin were incubated in the streptavidin coated channels for 30 mins in T100 buffer with 500 mM NaCl and 5 mM EDTA. The looping assay was performed in a reaction buffer (100 mM Tris-HCl pH 7.5, 100 mM NaCl, 7.5 mM MgCl2, 0.5 mg ml–1 BSA, 0.2 mM TCEP, 2 mM ATP, 50 nM SxO, 30 mM d-glucose, 2 mM trolox, 10 nM catalase, 37.5 µM glucose oxidase), and included 200 pM of labelled JetABC complexes, unless otherwise specified. The lower salt was used for promoting JetD association to JetABC (Supplementary Figure 1D). Cleavage assays with various DNA substrate were carried out using 200 to 300 pM concentrations of JetABCD and 5nM JetD in excess with 20 mM KCl or NaCl and 50 nM SxO in the reaction buffer. Whenever comparisons were made, identical buffer conditions and protein concentrations were used. These cleavage assays were performed at a lower 561 nm laser power (0.03 W/cm2) to minimize photodamage. For visualization with higher signal to noise ratio in Supplementary Movie 5, the laser power was doubled to 0.06 W/cm2.

### Data analysis

The image sequences stored were analyzed using a custom software previously developed in Python, available at https://github.com/biswajitSM/LEADS; (3), albeit with minor modifications. Through this software, regions containing the DNA substrates were cropped and saved in TIFF format. For quantitative analysis, the background noise was removed from the images using a ‘white_tophat’ filter, followed by smoothened with a median filter with a box size of 4 pixels (4). For the snapshots presented in the paper, the images with the DNA channel were only background subtracted whereas the images with the protein channel were both background subtracted and smoothened with the median filter. kymographs were constructed from the processed images by summing fluorescence intensities across 11 pixels centered on the DNA axis (e.g. Figure 1E). Each vertical line in the kymograph represents a single frame (time point) from the image sequence. Intensities in the kymograph were subtracted from the background intensities in the kymograph containing no DNA.

The kymographs were analyzed to obtain the sizes of DNA within the loop/punctum and in the regions outside the loops. The positions of the DNA puncta were obtained by finding local maxima (‘find_peaks’ in scipy) on each line of the kymograph. Subsequently, less prominent peaks with lower intensities were excluded through a manually set threshold. Typically, there is one punctum per line. The intensity withing the loop (*Intloop*) was calculated by summing the intensities of 9 pixels centered on the positions of the puncta. The remaining intensities on either side of the puncta were arbitrarily classified as up- and down-intensities respectively (*Intup* and *Intdown*). The fluorescence intensities in these distinct regions were then converted to DNA sizes in base pairs. This was achieved by dividing the fraction of intensities in each region by the total DNA size of 48.5 kb as indicated follows:

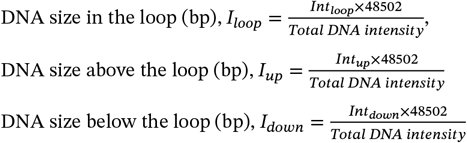

The sizes of DNA in various regions were plotted over time to analyze the kinetics of the loop or puncta formation. The rate of change in the size of the loop and up/down regions was determined by fitting the data with a linear equation (*I_x=rate×time+constant*, where x if loop, up, down) over a time window of 10 s, shifting this time window incrementally by one time point. This method yielded a rate vs. time (center of the time window) curve for each DNA molecule (Supplementary Figure 2D). The maximum rate from the looping rate vs. time data was designated as the loop extrusion rate (*rateloop*) for that specific DNA molecule. The up- and down-rate values corresponding to the time point of *rateloop* were noted as the respective up- and down-rates (*rate*_*up*_ and *rate*_*down*_) for that DNA molecule (Figure 1J and Figure 2J).

A scatter plot between rate_up_ and rate_down_ was utilized to assess the proportion of symmetric loop extrusion events (Figure 1K and 2K). If the values of both *rate*_*up*_ and *rate*_*down*_ were negative and less than -0.1 kbp/s, the event was classified as symmetrical loop extrusion.

The stalling force (e.g., Figure 1M) was determined based on the relative extension of the DNA when the loop reached its maximum size (Supplementary Figure 2F-G). The relative extension was calculated using the formula *l/(0.33nm/bp (Iup+Idown))*, where *l* represents the end-to-end distance of the double tethered DNA measured in nanometers. The stalling force was then deduced by interpolating the experimental data of relative-extension vs force, as reported by Ganji et. al.(5).

The loop’s pixel positions were used to extract the intensity of the protein signal from the kymograph of the protein channel. The number of fluorophores/labels (Figure 1G, 2I and 3G) was estimated from the intensity time trace. Events where the loop exhibited no protein signal were denoted as having 0 labels. When the loop extrusion events had clear bleaching (complete disappearance of the signal) that occurred in a stepwise manner, the number of labels were categorized as 1, 2, and >2, corresponding to the count of bleaching steps observed in the time trace. In the cases when no bleaching was seen within the observation time, the number of labels was estimated by comparing the intensities of those of nearby molecules in the imaging area that had undergone bleaching.

Protein signals not colocalized with any DNA loops were categorized as either translocation or immobile events (Figure 2L). Binding events were deemed immobile if the mean position remained constant over time. Conversely, they were considered as translocation events if the signal moved unidirectionally along the DNA.

### Plasmid cleavage analysis

Kymographs similar to those previously described were employed for plasmid cleavage analysis. An abrupt elongation of the DNA coupled with a sudden decrease in intensity was identified as a cleavage event (e.g., Figure 4A). When the intensity was uniformly distributed along the DNA’s length and no excess intensity was detected at the tether point, the event was classified as cleavage at the tether. Conversely, if additional intensity was detected near the tether point or if two distinct DNA strands became visible, the event was categorized a non-tethered cleavage (Figure 4C). In cases of non-tethered cleavage, the extra intensity was quantified and converted to the lengths expressed in kilobase pairs (Figure 4B) by assuming the total intensity corresponds to 44 kbp. Linear DNAs present at the start of the experiment were excluded from the cleavage analysis.

### Mass photometry

The mass photometry experiments were performed based on the protocol outlined in Pradhan et. al.(2) with slight modifications. A coverslip (1 mm thickness) was laser-drilled to make channels as previously detailed. Both the slide and coverslip were rinsed with water and 50% isopropanol alternatively two times and then dried using compressed air. Channels were constructed with double-sided adhesive tapes, and any opening other than the inlet and outlet were sealed using epoxy. The measurements were performed using a “TwoMP” instrument (Refyn Ltd.). The channels were equilibrated with MP buffer (50 mM HEPES pH 7.5, 20 mM KCl, 1 mM DTT, plus 10 mM MgCl2). The Wadjet samples were diluted in MP buffer before introduction into the flowcell. Imaging was done for 100 s with events being monitored in the TwoMP device. Initial analysis was done in the “DiscoverMP” software (Refyn Ltd.) and further quantifications were done in a custom written python script. For quantification of dimer and monomer fractions, landing events within a 200 to 1000 kDa range were considered. The area under the peaks were obtained from gaussian fitting and the fraction represented by the peak near 700 kDa was attributed to the dimer fraction (Figure 2B).

The error bars with 95% confidence interval in Figure 1G, 1K, 2D, 2K, 3G, 4D were calculated using the ‘binomial proportion confidence interval’. The error bar in Figure 2C, Supplementary Figure 1C shows the standard deviation values. Box whisker plots in Figure 1L-M, Figure 3K, Supplementary Figure 4D-E contain the respective median values (horizontal white line), with the box extending from Q1-Q3 quartile values of the data and the error bar extending no more than 1.5×IQR from the edges of the box.

