## Supplementary Material for "Loop extrusion-mediated plasmid DNA cleavage by the bacterial SMC Wadjet complex"

*The PDF file includes:*

Supplementary Figures 1-7

*Other Supplementary Materials for this manuscript include the following:*

Supplementary Movies 1 to 7

References

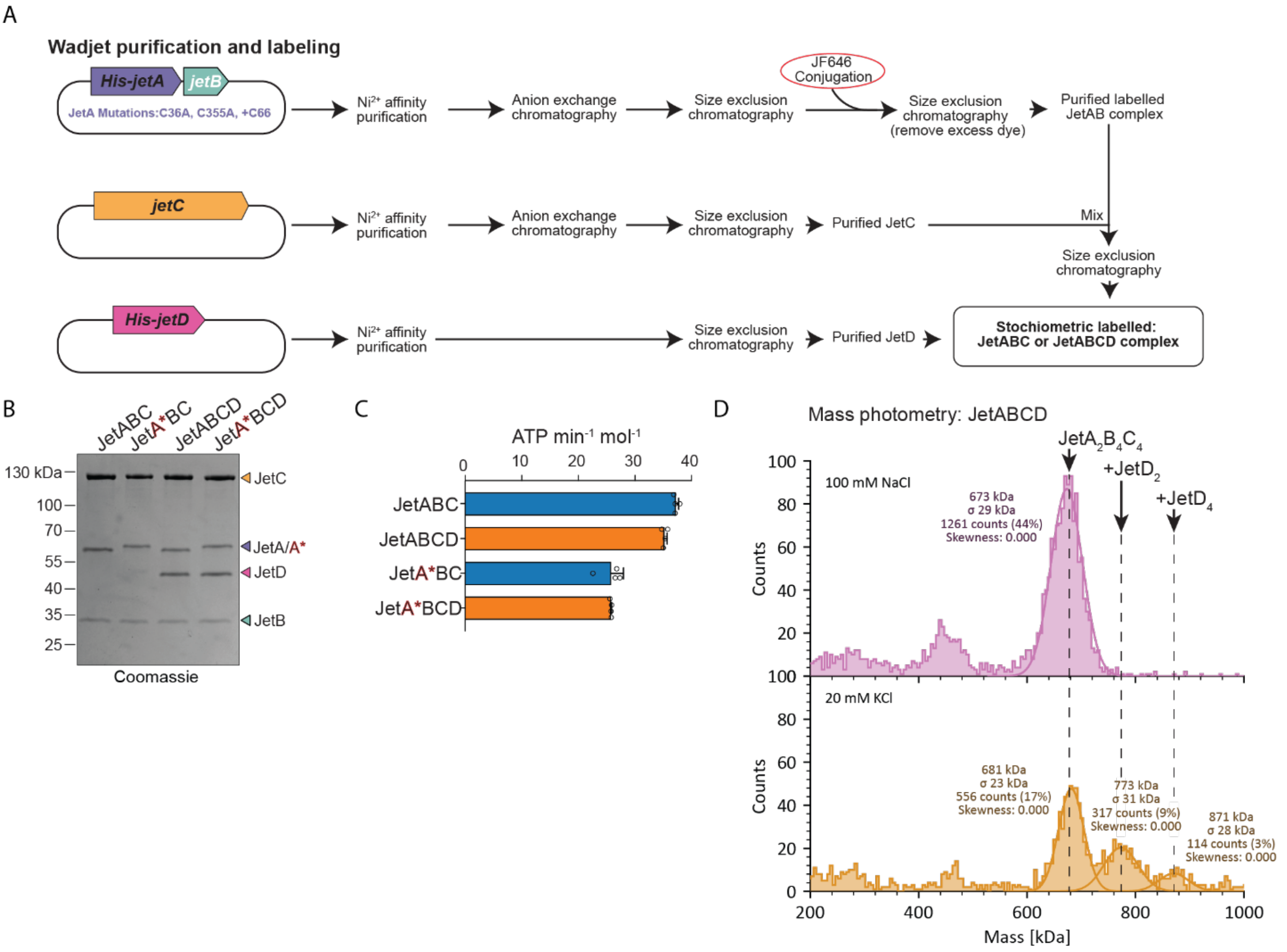


**Supplementary Figure 1. Purification, labeling, and characterization of Wadjet complexes. (A)** Purification and labeling scheme for *P. aeruginosa* PA14 Wadjet complexes. $His_6$-JetA (with C36A, C355A, mutations and C66 insertion) and JetB are coexpressed; untagged JetC is expressed on its own (untagged JetC binds strongly to a $Ni^{2+}$ affinity column), and $His_6$-JetD is expressed on its own. JetAB complexes are optionally incubated with Janelia Fluor 646 (JF646) dye, then mixed with JetC (yielding JetABC) or JetC and JetD (yielding JetABCD). **(B)** Coomassie blue-stained SDS-PAGE gel showing purified JetABC and JetABCD complexes. Red "A*" indicates that JetA has been JF646-labeled; labeling results in a small shift of the JetA band. **(C)** Mass photometry of JetABCD complexes in 100 mM NaCl buffer (top) or 20 mM NaCl buffer (bottom). In 100 mM NaCl buffer, JetD does not associate with JetABC. In 20 mM NaCl buffer, JetABC complexes bound to one or two dimers of JetD are observed. **(D)** ATP hydrolysis of JetABC and JetABCD complexes, shown as ATP molecules hydrolyzed per minute per full $JetA_2B_4C_4(D_4)$ complex. Data are graphed as the average +/- standard deviation of four independent measurements (datapoints shown as open circles). Red "A*" indicated that JetA has been JF646-labeled; labeling results in reduced ATPase activity.

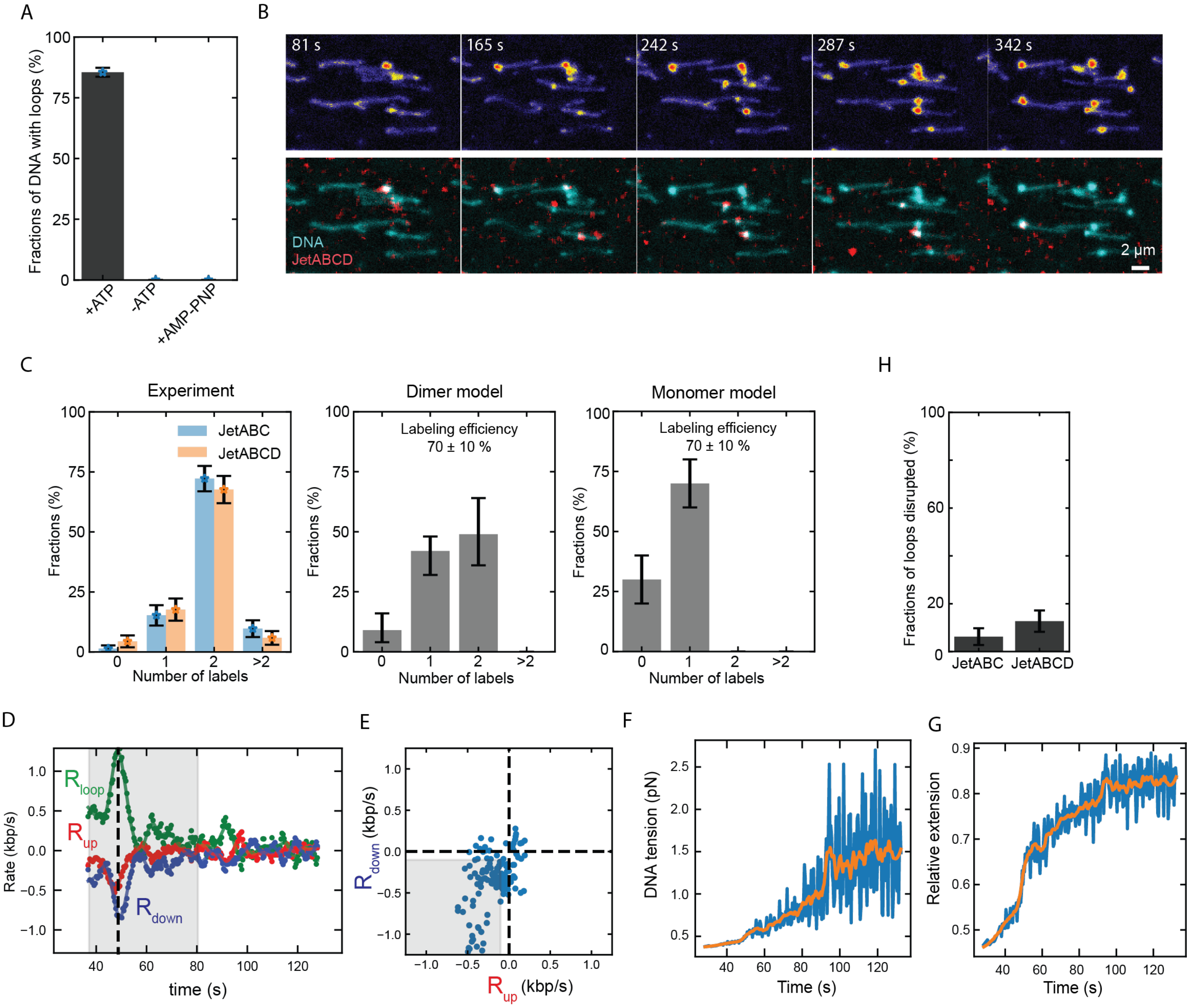


**Supplementary Figure 2. Single-molecule loop extrusion on double-tethered DNA. (A)** Fraction of double-tethered DNA undergoing loop extrusion by JetABC with or without ATP, or with AMP-PNP. N=366, 346, 332 for with ATP, without ATP, and with AMP-PNP, respectively. **(B)** Snapshots showing multiple (>5) double-tethered DNA molecules (top) and overlay with JetABC (bottom) at different time points following the addition of JetABC and ATP. **(C)** The experimental result (left, duplicate of Figure 1G) and the calculated probabilities (middle, right) for finding none, one or two labels assuming that all loop-extruding Wadjet complexes are dimers (middle) or monomers (right) with labelling efficiency of 70 ± 10%. The error bars in the models correspond to the standard deviation in the labelling efficiency. **(D)** The rates of the DNA length-changes within the loop ($R_{loop}$) and outside the loop ($R_{up}$, $R_{down}$) calculated from the DNA lengths shown in Figure 1I using a time window of 10 secs and shifting the window by one time point each time. The values of Rate up and Rate down in Figure 1J were taken at the time point where loop growth rate was maximal (dashed line). **(E)** Scatter plot showing the rate distributions $R_{up}$ and $R_{down}$ for the DNA molecule shown in (D). The displayed data points were taken from the grey shaded area in (D). **(F)** The relative extensions of the double-tethered DNA over time during loop extrusion and the corresponding **(G)** tension on the DNA, estimated from the known force-extension curve. The stalling force values in Figure 1M were taken from the value of the smoothened curve (orange) after the relative extension reached a plateau. **(H)** Fraction of DNA loops disrupted by JetABC and JetABCD during the measurement duration of 1000 seconds.

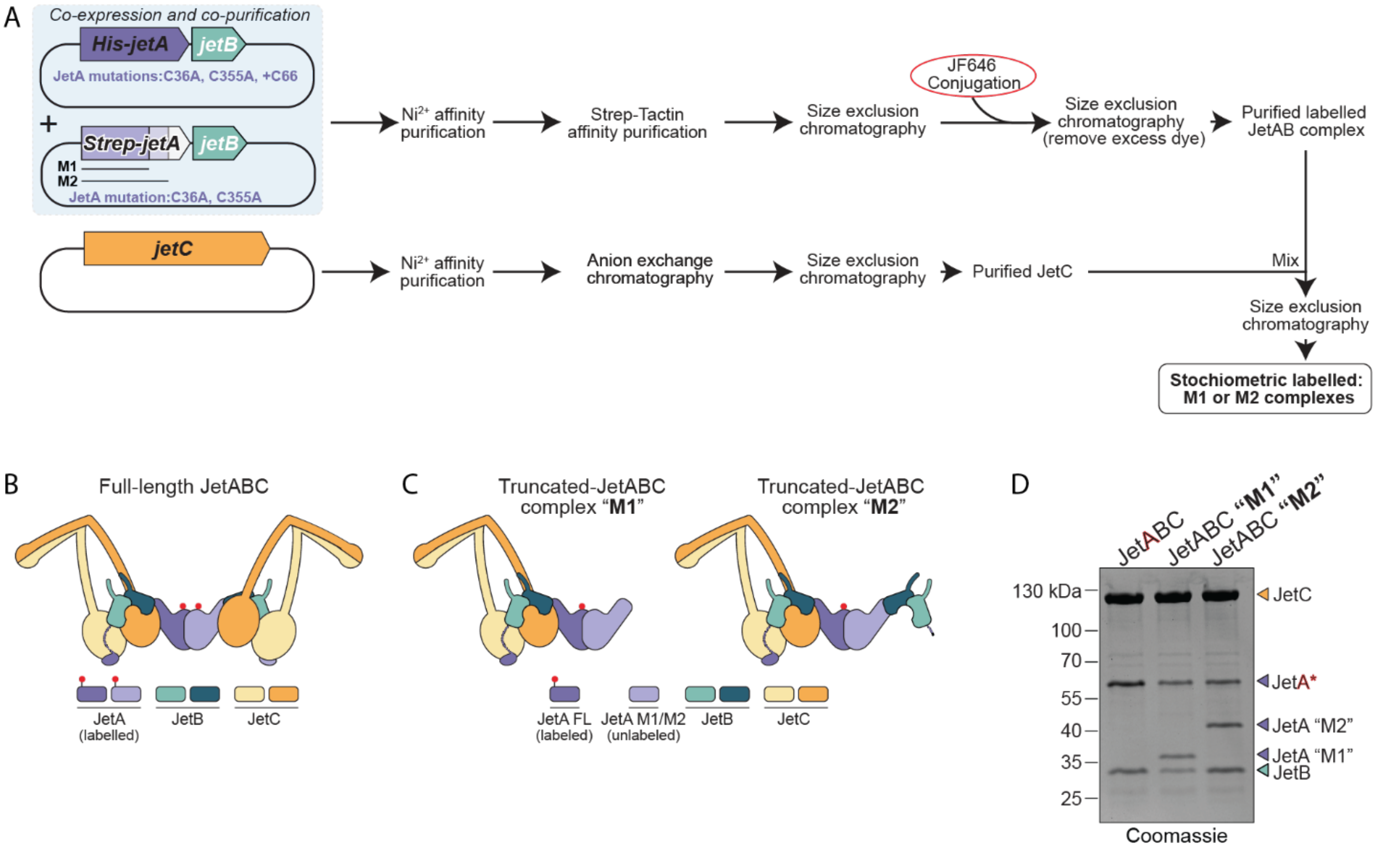


**Supplementary Figure 3. Purification of monomeric JetABC M1 and M2 complexes. (A)** Purification and labeling scheme for JetABC complexes with one full-length and one truncated JetA protomer. Full-length His$_6$-JetA (with C36A, C355A, and mutations and C66 insertion)+JetB is coexpressed with truncated (M1 or M2) StrepII-JetA (with C36A and C355A mutations)+JetB, then JetA heterodimers are purified by sequential $Ni^{2+}$ and Streptactin affinity chromatography. Complexes are then labeled with JF646 and mixed with JetC. **(B, C)** Schematics of full-length dimeric JetABC and the expected subunit composition of complexes carrying one truncated JetA subunit (M1 or M2). **(D)** Coomassie blue-stained SDS-PAGE gel showing purified JetABC complexes (full-length or carrying one truncated JetA subunit). Red "A" indicates that JetA has been JF646-labeled; labeling results in a small shift of the JetA band.

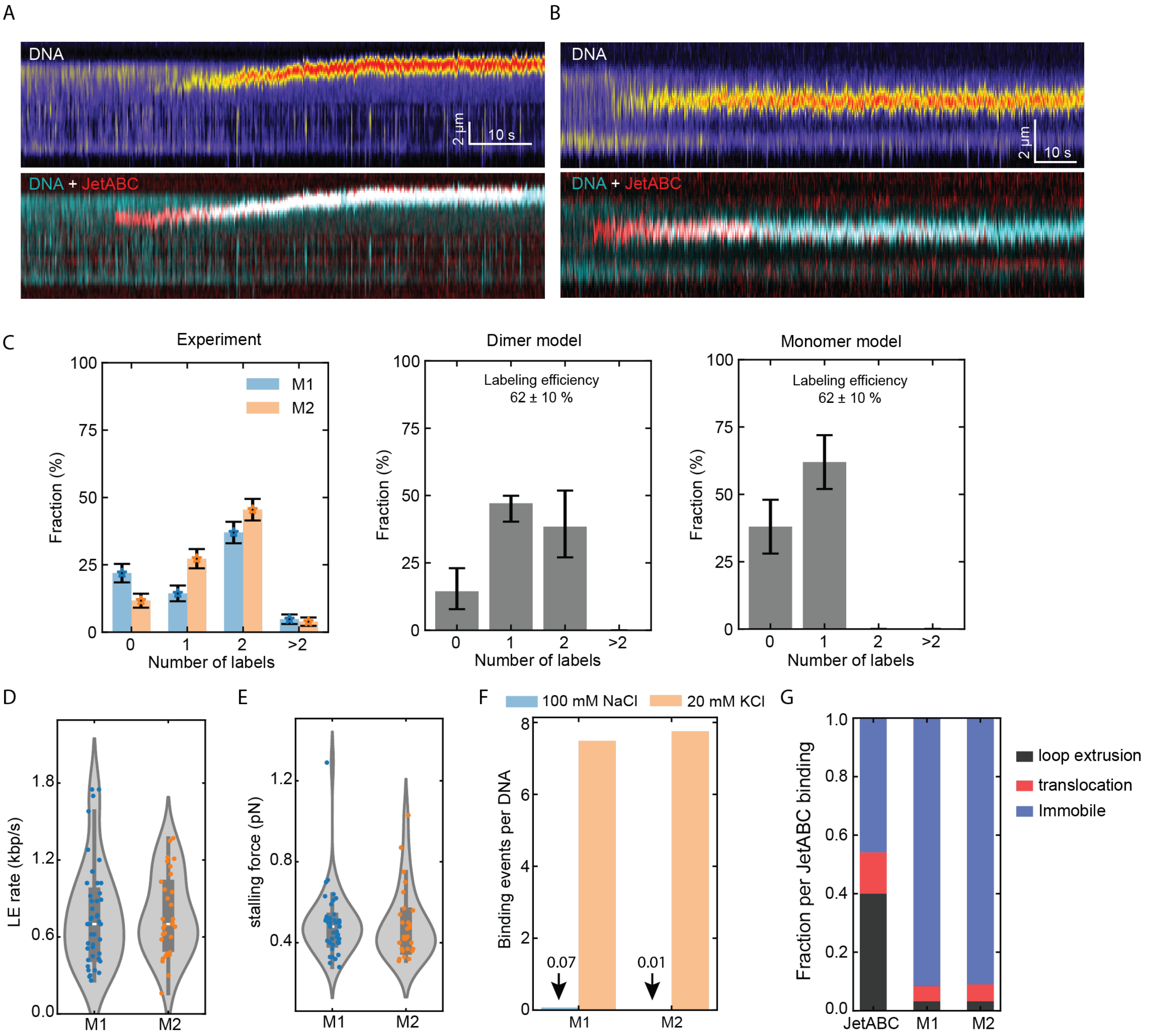


**Supplementary Figure 4. Loop extrusion by truncated JetABC. (A, B)** Kymographs of DNA (top) and overlay of DNA (cyan) and truncated-JetABC (red) (bottom) for M1 (A) and M2 (B) mutants. These kymographs were used for the plots shown in Figure 2E-H. **(C)** The experimental result (left, duplicate of Figure 2I) and the calculated probabilities (middle, right) for finding none, one or two labels assuming that all loop-extruding Wadjet complexes are dimers (middle) or monomers (right) with labelling efficiency of 62 ± 10%. The error bars in the models correspond to the standard deviation in the labelling efficiency. **(D)** Violin plot of loop growth rates for M1 (N=45) and M2 (N=31). **(E)** Violin plot showing the distribution of stalling forces for M1 (N=43) and M2 (N=28). **(F)** Comparison of number of M1 or M2 DNA binding events per DNA molecule for high salt buffer (100 mM NaCl) and low salt buffer (20 mM KCl). **(G)** Fraction of the Wadjet binding leading to DNA loop extrusion, translocation, and immobile events at low salt buffer (20mM KCl) for WT, M1and M2. Low salt buffer increases binding probability >100-fold and also increases the probabilities of non-loop extruding events (~90%).

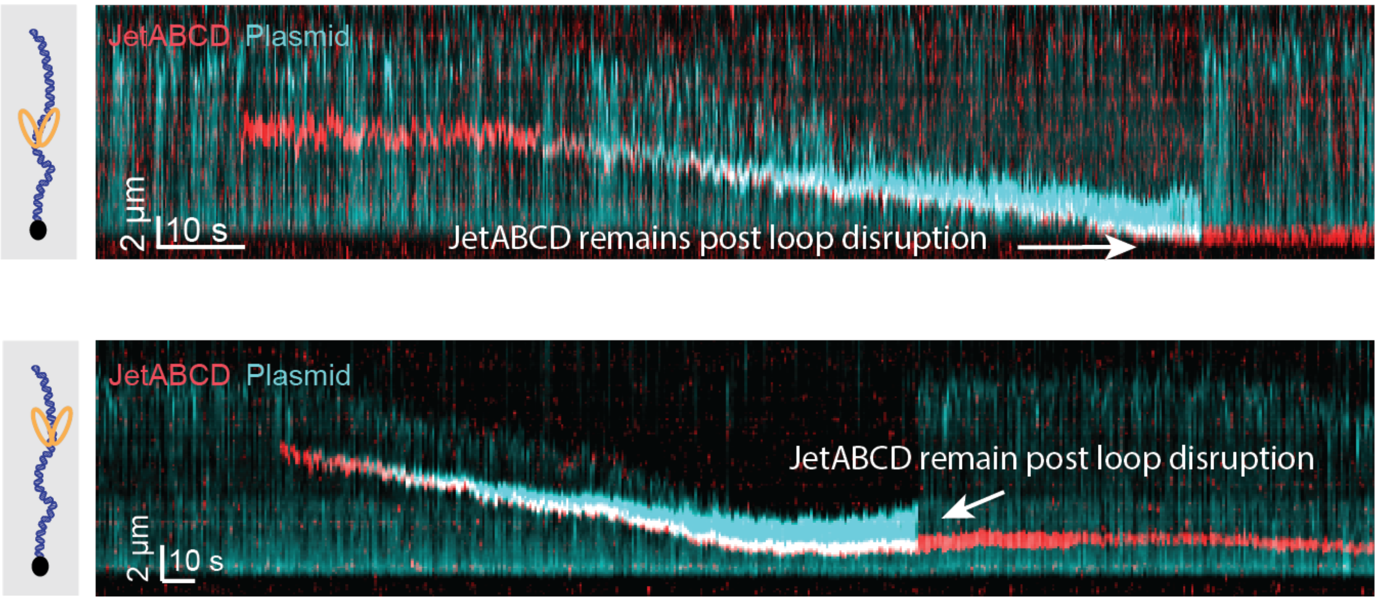


**Supplementary Figure 5. Accumulation of JetABCD complexes on linearized DNA.** Two overlay kymographs showing loop extrusion events on single-tethered DNA (cyan) leading to JetABCD (red) remaining near the tethered end of DNA.

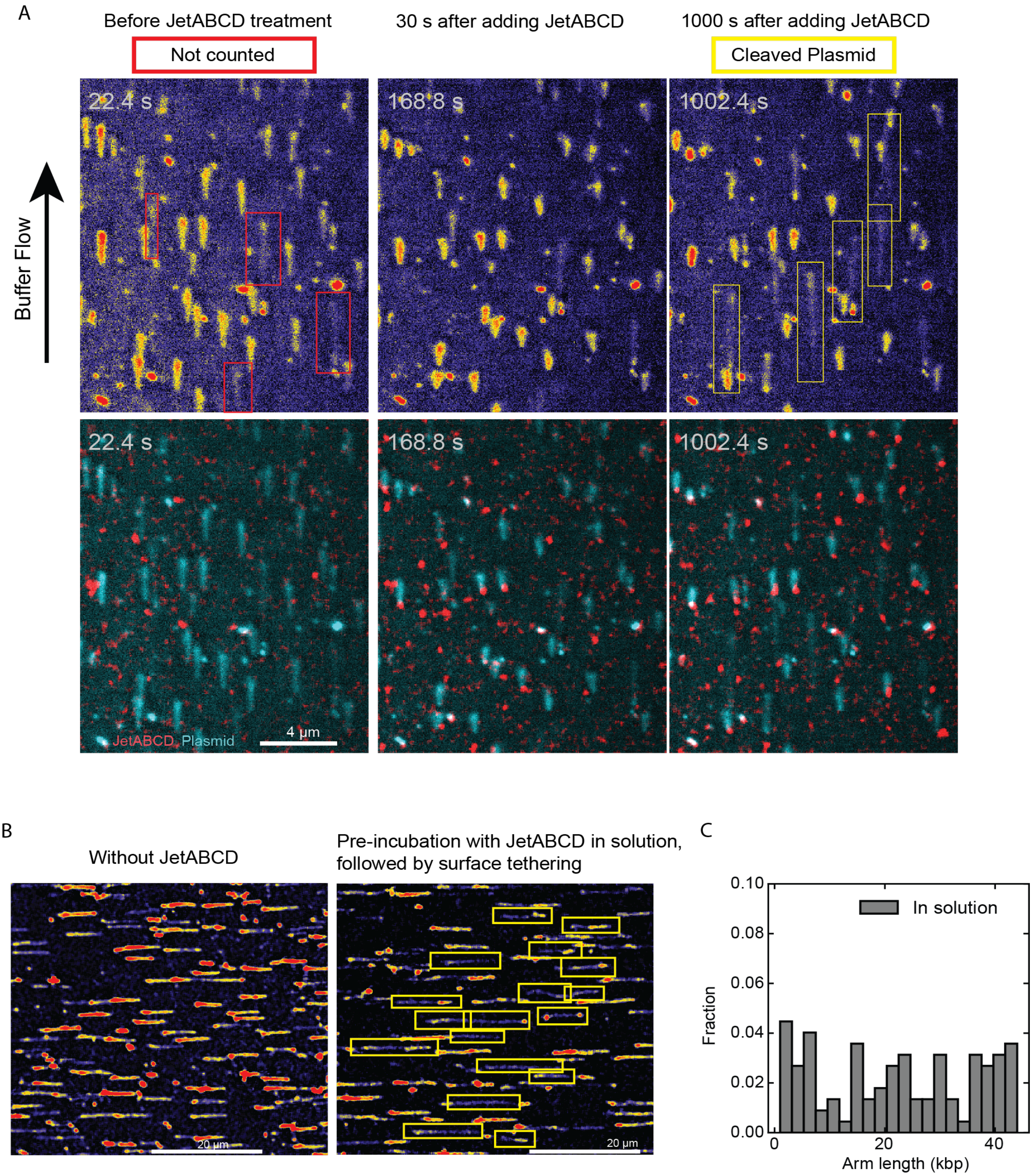


**Supplementary Figure 6. Comparison of plasmid cleavage by JetABCD for surface-anchored plasmids versus non-anchored plasmids in solution. (A)** Snapshots of DNA (top) and overlay with JetABCD (bottom) in a large field of view containing multiple tethered plasmids before and after treatment of 0.3 nM JetABCD and 5 nM JetD. Pre-existing linear DNAs (marked in red) were excluded from the analysis. The yellow boxes show the plasmids that were linearized after JetABCD-mediated loop extrusion. **(B)** Snapshots of DNA in a large field of view showing the products of plasmid cleavage (see Materials and Methods). In this case, the plasmids were pre-incubated with JetABCD in solution, followed by biotinylation of the cleaved products via Psoralen intercalation and the subsequent surface attachment of the flow cell surface. The yellow boxes indicate the cleaved DNA molecules. **(C)** Histograms showing the end-to-end length distributions of the buffer stretched-cleaved DNA as a result of the experiments in (B).

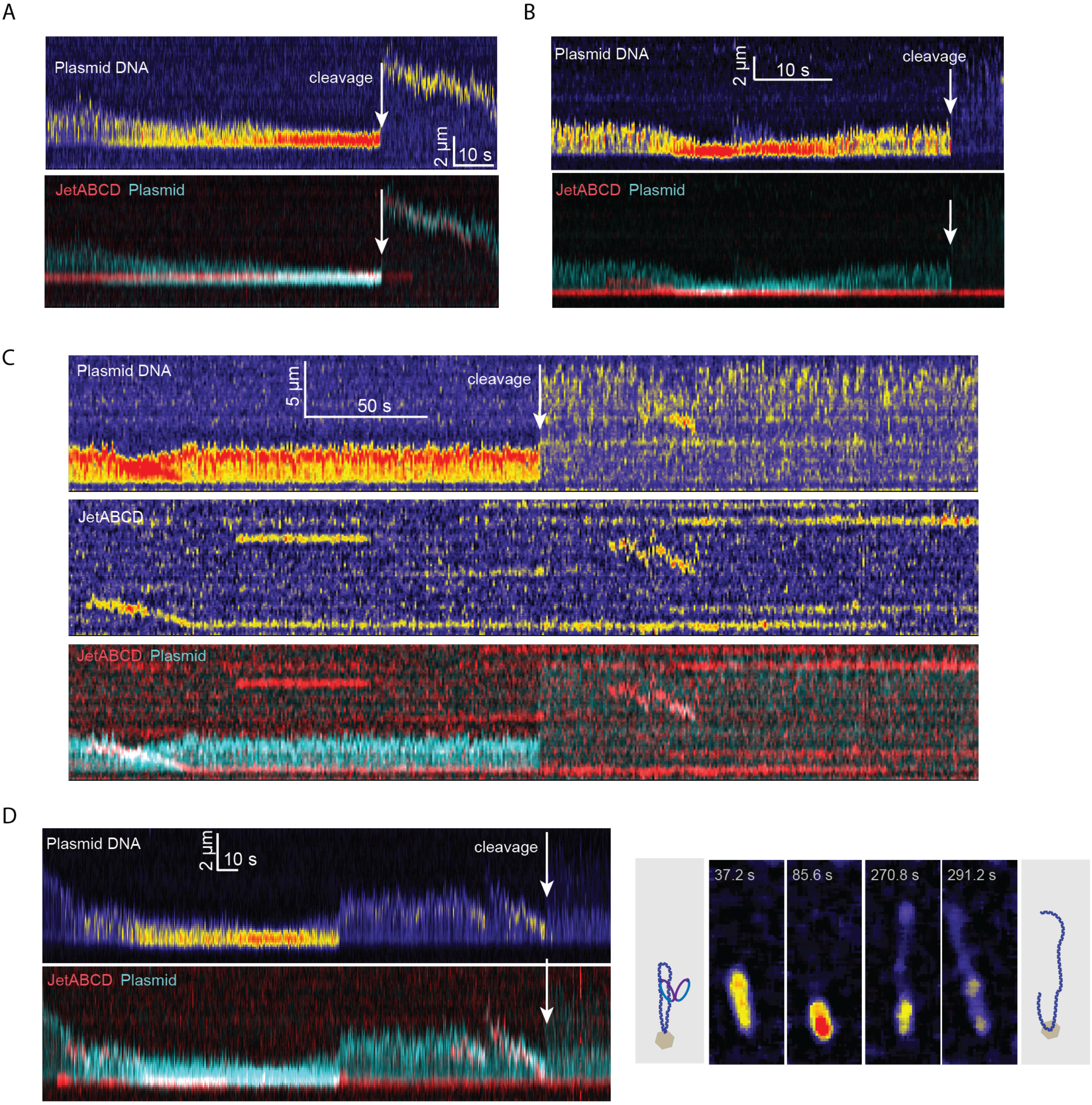


**Supplementary Figure 7. Additional examples of DNA Cleavage events followed by JetABCD-mediated loop extrusion and stalling. (A, B)** Two kymographs of DNA (top) and the overlay with JetABCD (bottom) showing the loop-extrusion mediated plasmid cleavage process, which occurred at the surface anchor position. **(C)** Single Wadjet loop extruding dimer complex leading to cleavage. The initial 120 seconds of this event was shown in Figure 4G. **(D)** Kymographs (left) and the snapshots (right) showing an example of plasmid cleavage that occurred at a position distant from the anchor point of the plasmid, as observed by the additional SxO-fluorescence intensity of the cleaved DNA near the tether point.

### Supplementary Movies 1-7

#### Supplementary Movie 1
DNA loop extrusion mediated by single fluorophore-labeled JetABC on double-tethered DNA under constant buffer flow.

#### Supplementary Movie 2
DNA loop extrusion by JetABC on double-tethered DNA without buffer flow.

#### Supplementary Movie 3
A large field of view movie showcasing examples of loop extrusion events by JetABC on multiple double-tethered DNA molecules.

#### Supplementary Movie 4
Multiple loop extrusion events performed by JetABC on surface-anchored plasmid DNA. Individual loop-extruding complexes stall upon reaching to the surface anchor points, in turn leading to accumulation of multiple complexes at the anchor points over time.

#### Supplementary Movie 5
A large field of view movie showcasing examples of plasmid DNA cleavage events occurred in multiple plasmids through loop-extruding JetABCD complexes.

#### Supplementary Movie 6
Plasmid DNA cleavage event occurred at the surface anchor point of the plasmid mediated by JetABCD after multiple loop extrusion events.

#### Supplementary Movie 7
Cleavage of a plasmid DNA following a single loop extrusion event by JetABCD.
